# AMPGLDA: Predicting LncRNA-Disease Associations Based on Adaptive Meta-Path Generation and Multi-Layer Perceptron

**DOI:** 10.1101/2024.06.02.596998

**Authors:** Dengju Yao, Xuehui Zhang, Xiaojuan Zhan

**Author notes:** Corresponding author. (Yao DJ). School of Computer Science and Technology, Harbin University of Science and Technology, Harbin 150080, China.

## Abstract

Increased evidence suggests that long non-coding RNA (lncRNA) holds a vital position in intricate human diseases. Nonetheless, the current pool of identified lncRNA linked to diseases remains restricted. Hence, the scientific community emphasizes the need for a reliable and cost-effective computational approach to predict the probable correlations between lncRNA and diseases. It would facilitate the exploration of the underlying mechanisms of lncRNA in ailments and the development of novel disease treatments. In this study, we propose a novel approach for predicting the associations between lncRNAs and diseases, which relies on the adaptive meta-path generation (AMPGLDA). Firstly, we integrate information about lncRNA, diseases, and miRNAs to construct a heterogeneous graph. Then, we utilize principal component analysis to extract global features from nodes. Based on this heterogeneous graph, AMPGLDA adaptively generates multiple meta-path graph structures and uses a graph convolutional neural network to learn the semantic feature representations of lncRNA and disease from the meta-path. Ultimately, AMPGLDA utilizes a deep neural network classifier to accurately predict the association between lncRNA and disease. The AMPGLDA model achieves impressive results, with AUC and AUPR scores of 99.66% and 99.66%, respectively, under the independent test set. Furthermore, three case studies demonstrate its accuracy in discovering new lncRNA-disease associations.

## Introduction

LncRNA, representing a class of non-coding RNA molecules, encompasses transcrips exceeding 200 nucleotides in length [1]. Increased studies have revealed that lncRNA not only performs specific functions in various biological processes and participates in cellular activities [2-5], but its abnormal expression is also often associated with the occurrence and progression of diseases [6-8]. Therefore, accurately identifying disease-related lncRNA is essential for exploring the etiology, development processes, and mechanisms of diseases. This information can assist in clinical diagnosis, disease prevention, and advancement of treatment.

Numerous research studies have devised various computational models to predict lncRNA-disease association (LDA), aiming to decrease the expense of discovering disease-related lncRNA via biological experimentation. The categorization of these LDA prediction models encompasses three primary groups.

The first group of LDA prediction models incorporates diverse biological information about lncRNA and integrates various biological networks to predict disease-related lncRNAs [9]. For instance, Ping et al. [10] combined two similarities of lncRNA with disease to calculate their association. Sun et al. [11] implemented a restarted random walk algorithm on the lncRNA functional similarity network, thereby enabling the anticipation of LDA. Nonetheless, this approach loses its utility for unknown disease candidate lncRNA. Furthermore, in the case of new lncRNA-related diseases, this method becomes less effective due to the absence of biological experimental validation. To address this issue, Chen [12] combined established connections between lncRNAs and diseases, lncRNA expression patterns, functional similarities among lncRNAs, semantic similarities related to diseases, and Gaussian interaction profile kernel similarities to unveil potential lncRNA-disease associations. While this model can predict lncRNA for unknown related diseases, it is impeded by limited tissue-specific expression information and low expression levels of lncRNAs.

Secondly, the LDA prediction methods based on machine learning have proven successful in tackling a range of prediction issues in the field of biology. These methods can use different datasets to foresee the potential association between lncRNA and disease, they have facilitated the discovery of actual lncRNA using further experiments. For example, Chen et al. [13] designed a prediction model based on the Laplacian regularized least squares method to predict LDAs (LRLSDA). However, to further improve LRLSDA, Chen et al. [14] presented two novel methods to calculate lncRNA functional similarity. Drawing upon incremental principal component analysis (PCA) and RF algorithm, Zhu et al. [15] proposed a novel integrated machine learning-based approach (IPCARF). Other methods include predicting lncRNA-disease associations through support vector machines (SVMs) [16], random forests (RF) [17-19], matrix factorization [20,21], random walks [22-24], and naive Bayesian classifiers [25].

Thirdly, deep learning-based LDA prediction techniques have garnered attention in recent years. For instance, graph attention networks have been applied to forecast LDA. Lan et al. [26] proposed GANNLDA, leveraging a graph attention network to extract pertinent features from the characteristics of lncRNA and disease. Subsequently, they employed a multi-layer perceptron (MLP) to predict lncRNA-disease associations. Furthermore, novel strategies have emerged, such as Xuan et al. [27] introduced a neighbor topology coding, which facilitates the learning of the neighbor topology of nodes in the heterogeneous graph. Models incorporating convolutional neural networks have also been developed for predicting lncRNA-disease associations [28–30]. As an instance, Xuan et al. [31] introduced ACLDA, a method designed to deduce potential lncRNA candidates linked to diseases, ACLDA integrates a fully connected autoencoder and convolutional neural network. Zhao et al. [32] conceptualized an innovative framework that leverages a heterogeneous graph attention network grounded in the utilization of meta-paths. Xuan et al. [33] integrated lncRNA and disease properties with meta-path semantics to predict disease-associated ncRNAs, enhancing prediction accuracy. Sheng et al. [34] utilizes graph contrast learning to extract latent topological features from nodes.

While the existing approaches have demonstrated satisfactory predictive performance, these methods are computationally complex, and there are still some shortcomings in these models. Notably, most researchers have not fully noticed the wealth of semantic information within the lncRNA-disease heterogeneous graph. To address this limitation and better harness the information contained in heterogeneous graphs, we consider combining neural networks with meta-paths to consolidate node feature information and meta-path information within the heterogeneous graph,thereby enhancing overall model performance. Meta-paths, representing paths connecting different types of nodes, serve as a valuable tool for extracting complex structural and rich information within heterogeneous networks. Currently, only a handful of meta-path-based association prediction models rely on researchers’ biological knowledge to define meta-paths, potentially introducing noise and redundancy and compromising the model performance. To overcome these challenges and to more fully exploit the potential of meta-path research, our study proposes a cutting-edge lncRNA-disease association prediction model, named AMPGLDA, which is underpinned by an adaptively generated meta-path, offering a novel perspective in this field. The principal contributions of our model can be outlined below: (1) our heterogeneous graph accounts for the diversity of nodes and edge relationship types, utilizing several interconnected edges to capture the low-dimensional features that are to each individual node. We introduce miRNA biological information and use the association, similarity, and interaction information between lncRNA, disease, and miRNA to construct a three-layer heterogeneous graph with diverse node and edge types, providing a comprehensive depiction of relationships between nodes; (2) our model benefits from the adaptive generation of meta-paths, eliminating the dependence on biological expertise and allowing for the generation of various meta-paths of any length. This approach reduces redundancy and noise that may arise from the manual meta-path definition, ultimately enhancing model performance; (3) the PCA dimensionality reduction method and graph convolutional neural network are employed respectively to explore the global and semantic feature representation of lncRNA and disease. A comprehensive evaluation including ablation experiments, comparison with recently published models, case studies, survival analysis of breast cancer, and experimental analysis of multiple datasets, demonstrates the good predictive performance and strong generalization ability of our model in disease-related lncRNA prediction.

## Results

In this section, we initially elucidate our evaluation metrics. To gauge the robustness of our model, AMPGLDA underwent comprehensive experimental analyses across various datasets within the same experimental environment, comparing its performance with several other state-of-the-art LDA prediction models.

### Evaluation metric

To assess the predictive performance of AMPGLDA, we utilize 5-fold cross-validation, which enables us to ascertain its efficacy in preventing overfitting and the credibility of the model’s generalization capabilities. For the construction of the positive and negative sample set, we designate unverified associations as negative samples, while all authenticated lncRNA-disease associations constitute the positive samples. Subsequently, we randomly select an equal number of both sample types to form the final dataset, which is then divided randomly into 80% training samples and 20% independent test samples. Within the cross-validation process, the training samples are randomly partitioned into five equally sized subsets. One of which in turn served as a test set and the other four as a training set, and repeated the training and testing five times, to fully evaluate the performance of the model and reduce the impact of randomness on the evaluation results. Finally, the model is applied to the independent test set which accounts for 20% of the dataset, and the test results are obtained. Primary evaluation metrics encompass the area under the ROC curve (AUC) and the area under the Precision-Recall curve (AUPR). Additionally, we calculate ACC, Precision, Sensitivity, F1-score, and MCC to provide a comprehensive assessment of our model’s performance.

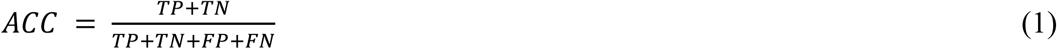

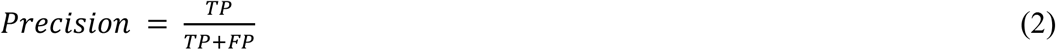

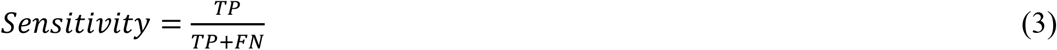

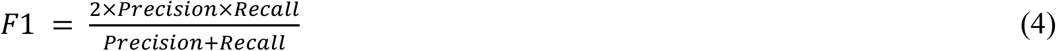

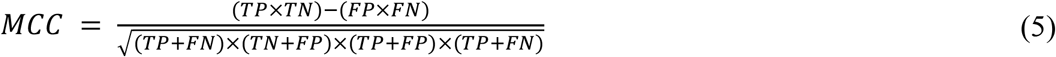

Here, TP (TN) signifies a true positive (true negative) instance, while FP (FN) denotes a false positive (false negative) instance.

### Comparison of performance evaluation between MLP and alternative classifiers

In the AMPGLDA model, we used the multi-layer perceptron (MLP) classifier for predicting lncRNA-disease associations. To confirm the suitability of MLP, we conducted a comparative analysis of its performance against several classical classifiers, including RF, SVM, eXtreme Gradient Boosting (XGBoost), Stacking Ensemble Classifier (Stacking), Gradient Boosting Decision Tree (GBDT), and based on a common dataset. Under 5-fold cross-validation, we evaluated the performance of these six classifiers based on metrics such as AUC, AUPR, Precision, Sensitivity, F1-score, ACC, and MCC. **Table 1** presents the results of these evaluations. Our analysis indicates that MLP outperforms other classifiers across all performance metrics, attaining the highest values for AUC, AUPR, Precision, Sensitivity, F1-score, ACC, and MCC, at 99.66%, 99.66%, 97.68%, 98.54%, 98.11%, 98.07%, and 96.15%, respectively. Consequently, we have chosen MLP to predict the associations between lncRNAs and diseases.

**Table 1.** The performance comparison between MLP and other classifiers.

| Method | AUC | AUPR | Precision | Sensitivity | F1 | ACC | MCC |
| --- | --- | --- | --- | --- | --- | --- | --- |
| RF | 0.9948±0.0014 | 0.9947±0.0013 | 0.9681±0.0077 | 0.9763±0.0067 | 0.9722±0.0054 | 0.9726±0.0049 | 0.9452±0.0097 |
| GBDT | 0.9869±0.0026 | 0.9841±0.0044 | 0.9318±0.0050 | 0.9493±0.0074 | 0.9405±0.0051 | 0.9416±0.0048 | 0.8833±0.0096 |
| XGBT | 0.9936±0.0011 | 0.9932±0.0014 | 0.9691±0.0068 | 0.9854±0.0059 | 0.9772±0.0055 | 0.9772±0.0052 | 0.9545±0.0104 |
| SVM | 0.9648±0.0054 | 0.9627±0.0082 | 0.8931±0.0134 | 0.9053±0.0067 | 0.8991±0.0085 | 0.8971±0.0076 | 0.7943±0.0150 |
| Stacking | 0.9933±0.0017 | 0.9901±0.0021 | 0.9663±0.0052 | 0.9879±0.0041 | 0.9770±0.0040 | 0.9766±0.0040 | 0.9535±0.0080 |
| MLP | **0.9966±0.0010** | **0.9966±0.0012** | **0.9768±0.0071** | **0.9854±0.0064** | **0.9811±0.0037** | **0.9807±0.0039** | **0.9615±0.0078** |

The bold values indicate the higher metrics.

### Feature dimension analysis

Subsequently, we conducted a feature dimension analysis and determined the appropriate dimensions based on experimental results. Specifically, we compressed the dimensions of global features and semantic features to 16, 32, 64, 128, and 256, respectively. To eliminate the effect of randomness, we set the same random seed for MLP, ensuring more reliable and stable experimental results. The feature dimension that yielded the highest prediction performance was determined through experimental analysis. **Figure 1** illustrates the AUC and AUPR obtained from different dimensional features on the test set. The figure indicates that the maximum AUC and AUPR are attained when 128 dimensions are selected as global and semantic features. Therefore, 256 dimensions were chosen as the final feature dimensions.

**Figure 1.**
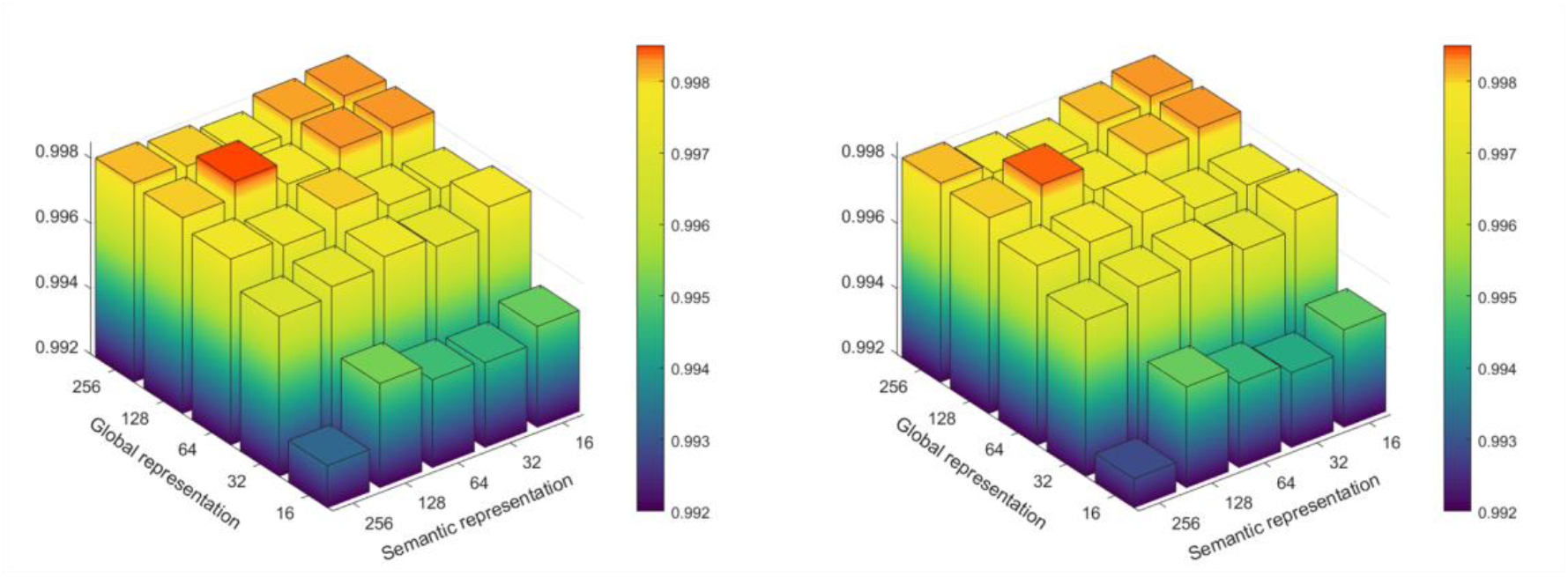
The AUCs and AUPRs under different feature dimensions Ablation study

In ablation experiments, we evaluated the impact of specific components or features on the overall performance of the model by purposely excluding them. In this particular investigation, our focus centered on features extracted from PCA (AMPGLDA-P) and those acquired through meta-path learning (AMPGLDA-M). Employing the AMPGLDA model as the baseline, we conducted a comparative analysis between features extracted solely from PCA and those obtained exclusively through meta-path learning, utilizing a 5-fold cross-validation. The experimental results are detailed in **Table 2**. In the case of using the meta-path method AMPGLDA-M, AUC and AUPR reached 96.90% and 96.63%, respectively. In the case of using only the PCA method AMPGLDA-P, AUC and AUPR reached 98.83% and 98.70%, respectively. In the final complete model AMPGLDA, compared to AMPGLDA-M and AMPGLDA-P, AUC increased by 2.76% and 0.83%, respectively, and AUPR increased by 3.03% and 0.96%. Ablation experiments substantiate the assertion that the synergistic integration of PCA and meta-path learning is both rational and efficacious. demonstrate the effectiveness and contribution of PCA and meta-path learning.

**Table 2.**
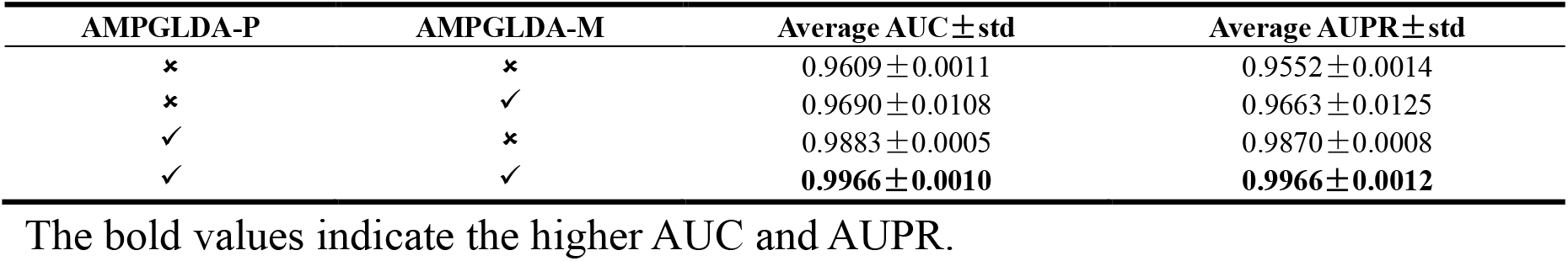
Ablation study results on AMPGLDA.

| AMPGLDA-P | AMPGLDA-M | Average AUC ± std | Average AUPR ± std |
| --- | --- | --- | --- |
| × | × | 0.9609 ± 0.0011 | 0.9552 ± 0.0014 |
| × | ✓ | 0.9690 ± 0.0108 | 0.9663 ± 0.0125 |
| ✓ | × | 0.9883 ± 0.0005 | 0.9870 ± 0.0008 |
| ✓ | ✓ | **0.9966 ± 0.0010** | **0.9966 ± 0.0012** |

The bold values indicate the higher AUC and AUPR.

The experimental findings highlight the predominant contribution of the similarity feature extracted by PCA. One plausible explanation is that node similarity features encompass not only direct similarity associations between lncRNAs and diseases, but also similarity associations involving their adjacent lncRNAs and diseases. In contrast, the contribution of meta-path-based feature extraction to the results is secondary. When compared to node similarity, the acquisition of features based on meta-paths can be perceived as establishing an indirect relationship between lncRNAs and diseases, concealed within the structure of heterogeneous maps. Nevertheless, it constitutes a crucial component of the prediction model, capable of enhancing overall prediction performance.

### Comparison with other methods

To validate the effectiveness of our approach, we conducted a comparative analysis against the following state-of-the-art methods. This study involves running both the comparison method and the AMPGLDA model on an identical dataset, thereby ensuring a fair and thorough assessment.

IPCARF [15]: IPCARF introduces an innovative algorithm, rooted in the integration of machine learning technology. The algorithm integrates multiple similarity matrices and employs incremental principal component analysis (IPCA) to reduce their dimensionality. Ultimately, random forest is applied for predicting lncRNA-disease associations.

GCLMTP [34]: Implementing graph contrast learning, this method serves as an unsupervised embedding model to extract potential topological features of lncRNAs, miRNAs, and diseases from heterogenous graph. The model achieves this by maximizing the mutual information between the block representation of heterogenous graph and the corresponding high-level abstracts through the graph convolution network architecture.

MAGCNSE [35]: This approach embraces deep learning methodologies, leveraging Graph Convolutional Networks (GCN) and attention mechanisms to integrate multi-view data. Additionally, a Convolutional Neural Network (CNN) is employed to acquire low-dimensional representations for both lncRNAs and diseases. Ultimately, the integrated classifier is utilized to predict lncRNA-disease associations.

HOPEXGB [36]: HOPEXGB is a prediction model for recognizing disease-associated lncRNA and miRNA by high-order neighborhood-preserving embedding (HOPE) and XGBoost.

LDAformer [37]: LDAformer designs multi-hop topological feature extraction methods, and uses a Transformer encoder to predict disease-related lncRNA.

Here, we employed 5 rounds of 5-fold cross-validation to evaluate the performance of AMPGLDA in comparison with other advanced methods. **Table 3** demonstrates that, under the same experimental environment and data conditions, the AUC of AMPGLDA reached 99.66%, surpassing IPCARF by 3.65%, GCLMTP by 0.48%, MAGCNSE by 4.08%, HOPEXGB by 9.78%, and LDAformer by 1.79%. Additionally, the AUPR of AMPGLDA was 99.66%, surpassing IPCARF by 4.13%, 0.52% higher than GCLMTP, 6.06% higher than MAGCNSE, 22.99% higher than HOPEXGB, and 1.96% higher than LDAformer. These results indicate that our AMPGLDA model exhibits superior predictive power for lncRNA-disease associations.

**Table 3.** The performance comparison of different LDA prediction models.

| Method | AUC | AUPR | Precision | Sensitivity | F1 | ACC | MCC |
| --- | --- | --- | --- | --- | --- | --- | --- |
| AMPGLDA | **0.9966±0.0010** | **0.9966±0.0012** | **0.9768±0.0071** | **0.9854±0.0064** | **0.9811±0.0037** | 0.9807±0.0039 | **0.9615±0.0078** |
| IPCARF | 0.9601±0.0228 | 0.9553±0.0261 | 0.8850±0.0530 | 0.8903±0.0601 | 0.8875±0.0562 | 0.8882±0.0552 | 0.7764±0.1106 |
| GCLMTP | 0.9918±0.0022 | 0.9914±0.0027 | 0.9637±0.0028 | 0.9596±0.0105 | 0.9616±0.0051 | 0.9609±0.0054 | 0.9219±0.0108 |
| MAGCNSE | 0.9558±0.0004 | 0.9360±0.0007 | 0.9511±0.0009 | 0.9349±0.0009 | 0.9430±0.0005 | 0.8861±0.0011 | 0.9511±0.0009 |
| HOPEXGB | 0.8988±0.0027 | 0.7667±0.0049 | 0.9531±0.0042 | 0.7987±0.0055 | 0.8691±0.0030 | **0.9934±0.0001** | 0.8693±0.0029 |
| LDAformer | 0.9787±0.0156 | 0.9770±0.0153 | 0.9002±0.0328 | 0.9540±0.0436 | 0.9262±0.0363 | 0.9242±0.0368 | 0.8502±0.0741 |

The bold values indicate the higher metrics.

Subsequently, we employed paired tests to analyze differences between AMPGLDA and other comparative methods. **Table 4** illustrates that AMPGLDA outperformed the most advanced methods, achieving higher scores in both AUC and AUPR.

**Table 4.** Difference between AMPGLDA and benchmark methods tested by Paired t-test in terms of AUC and AUPR.

| Methods | IPCARF | GCLMTP | MAGCNSE | HOPEXGB | LDAformer |
| --- | --- | --- | --- | --- | --- |
| p-value of AUC | 7.278e-3 | 2.255e-3 | 3.83216e-13 | 1.02974e-12 | 3.3813e-2 |
| p-value of AUPR | 7.739e-3 | 4.508e-3 | 1.0918e-13 | 9.52755e-14 | 2.1337e-2 |

### Robustness testing

To verify the robustness of our AMPGLDA model, we applied our model to various datasets while maintaining the model parameters constant. Dataset 1 is the dataset used in this study, while Dataset 2 and Dataset 3 were obtained from LDAformer [37], and SVDNVLDA [38], respectively. Specifically, Dataset 2 consisted of 665 lncRNAs, 316 diseases, and 295 miRNAs, featuring 3383 LDAs, 8540 MDAs, and 2108 LMIs. Dataset 3 encompassed 861 lncRNAs, 431 diseases, and 437 miRNAs, with 4518 DAs, 4189 MDAs, and 8172 LMIs. The AUC values of our proposed model under 5-fold cross-validation were 0.9928, 0.9651, and 0.9718 for Datasets 1, Dataset 2, and Dataset 3, respectively. The AUPR values were 0.9920, 0.9613, and 0.9712 for Dataset 1, Dataset 2, and Dataset 3, respectively. More detailed performance metrics, including ACC, Precision, Sensitivity, F1-score, and MCC values in addition to AUC and AUPR, are listed in **Table 5**.

**Table 5.**
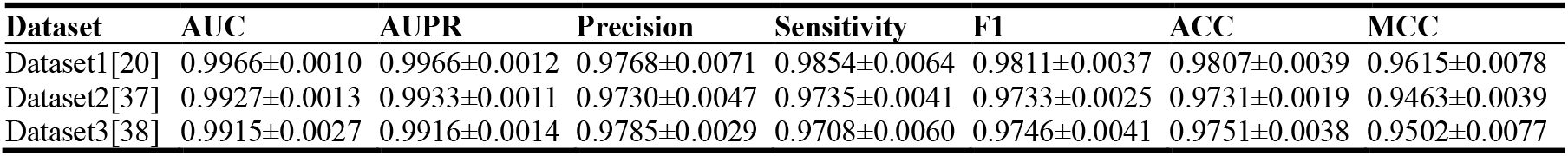
Experimental results across various datasets.

The results demonstrate that AMPGLDA exhibits robust predictive performance on Dataset 1, Dataset 2, and Dataset 3 under the same parameters. Hence, we conclude that the model showcases strong robustness across diverse datasets, indicating excellent generalization ability.

### Cases studies

In addition, through case studies of the three most prevalent cancers, namely breast, lung, and colorectal cancers, we explored the AMPGLDA model’s capacity for prediction. For each disease, known lncRNA-disease associations used in predictive models were excluded, and lncRNA candidates associated with these cancers were ranked based on their AMPGLDA scores, identifying the top 30 candidates for each cancer (listed in **Table 6, Table 7**, and **Table 8**, respectively). Moreover, we validated the predictions by checking public databases such as Lnc2Cancer v3.0, LncRNADisease v2.0, and the latest research literature to confirm associations with breast cancer, lung cancer, and colorectal cancers. In the evidence column of the table, “C” represents lncRNAs that have been validated as candidates by the Lnc2Cancer database, while “D” signifies candidate lncRNAs that are supported by the LncRNADisease database, “P” indicates that there is no evidence in the database but that the literature sources support it, and “N” means there is no evidence to prove it.

**Table 6.**
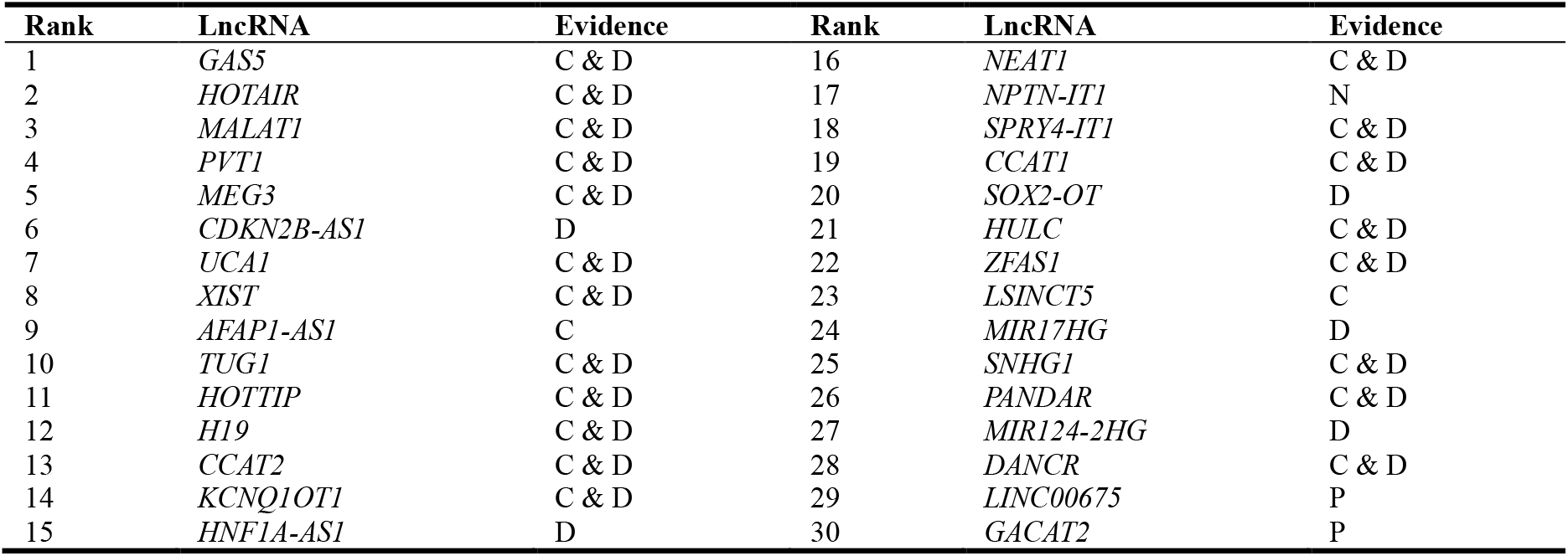
Top 30 lncRNA related to breast cancer predicted by AMPGLDA.

**Table 7.**
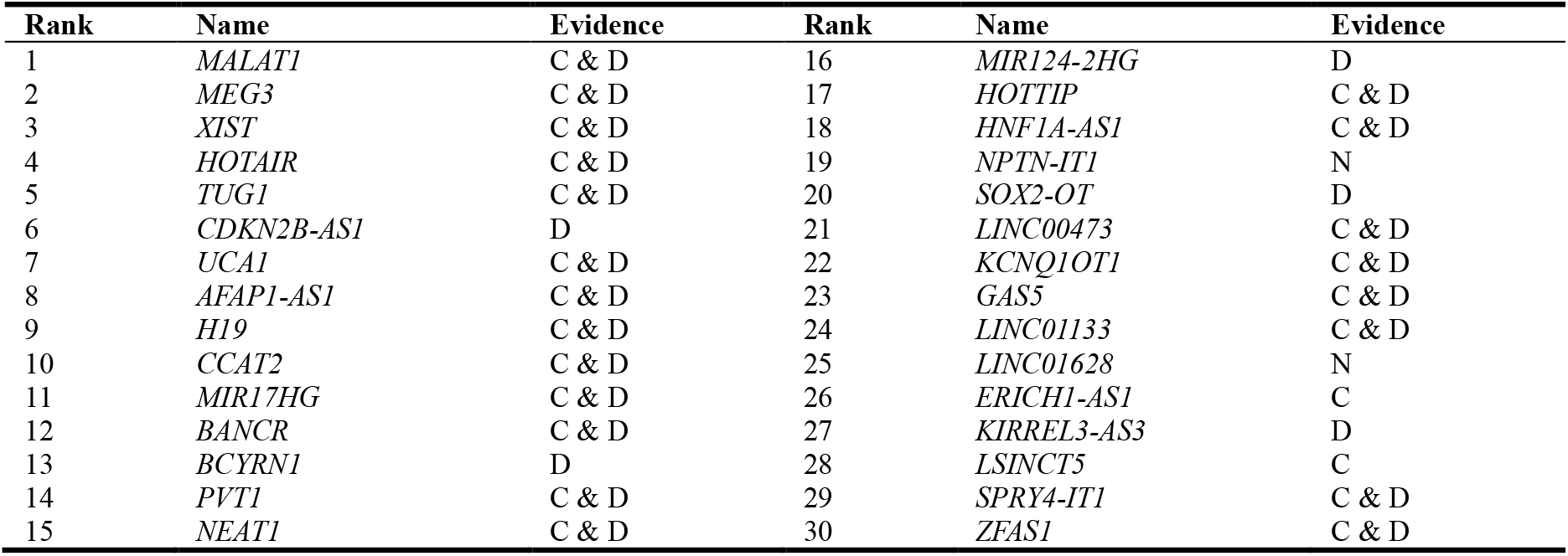
Top 30 lncRNA related to lung cancer predicted by AMPGLDA.

**Table 8.**
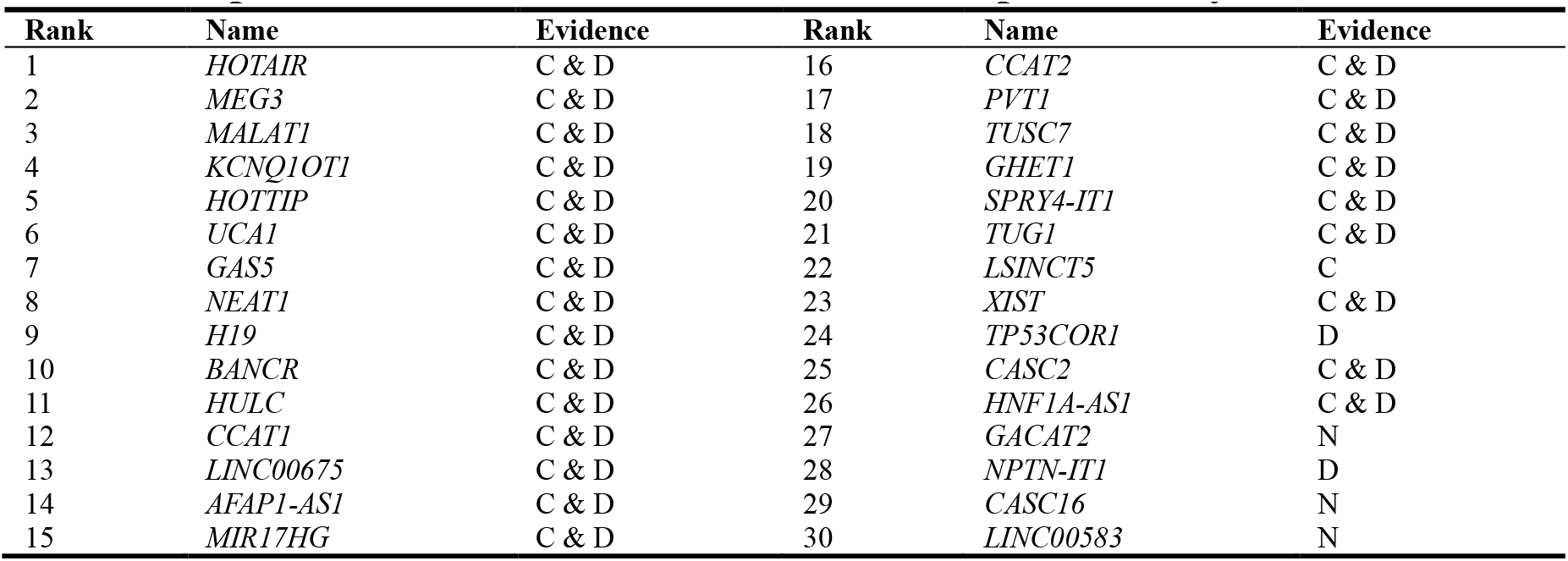
Top 30 lncRNA related to colorectal cancer predicted by AMPGLDA.

Of the top 30 lncRNAs listed in Table 6 that are most closely associated with breast cancer, 22 were also recorded in the Lnc2Cancer 3.0 database, confirming their association with the disease. Additionally, the LncRNADisease v2.0 database contains 25 candidate lncRNAs that were found to be abnormally expressed in breast cancer tissues. Recent research [39] has shown that *LINC00675* overexpression can inhibit breast cancer cell proliferation, migration, and invasion, while depletion of *LINC00675* has the opposite effect. Additionally, literature [40] has demonstrated that *GACAT2* is associated with breast cancer. Only one lncRNA candidate is labeled as ‘Unknown’, indicating a temporary lack of evidence supporting its association with breast cancer.

For lung cancer, 23 of the top 30 candidate lncRNAs listed in Table 7 were confirmed by the Lnc2Cancer 3.0 database, and 26 were confirmed by the LncRNADisease v2.0 database.

Table 8 lists the top 30 candidate lncRNAs related to colorectal cancer, with 25 verified by LncRNADisease and 26 by Lnc2Cancer. The ability of AMPGLDA to predict LDAs was further substantiated through these case studies involving these three cancers.

Furthermore, we selected HOTAIR and SOX2-OT lncRNA among the top 30 lncRNA candidates and performed survival analysis on these two lncRNAs predicted to be highly associated with breast cancer using TCGA data [41]. **Figure 2** was generated using online tools [42]. As depicted in Figure 2, over time, the expression of the lncRNA associated with breast cancer gradually decreases, and it correlates with an increase in survival time and probability for patients. The turning point for *HOTAIR* lncRNA is observed to lag behind that of *SOX2-OT*, indicating a greater likelihood of *HOTAIR*’s association with the disease. This observation suggests a positive prognostic effect of the predicted lncRNA on breast cancer.

**Figure 2.**
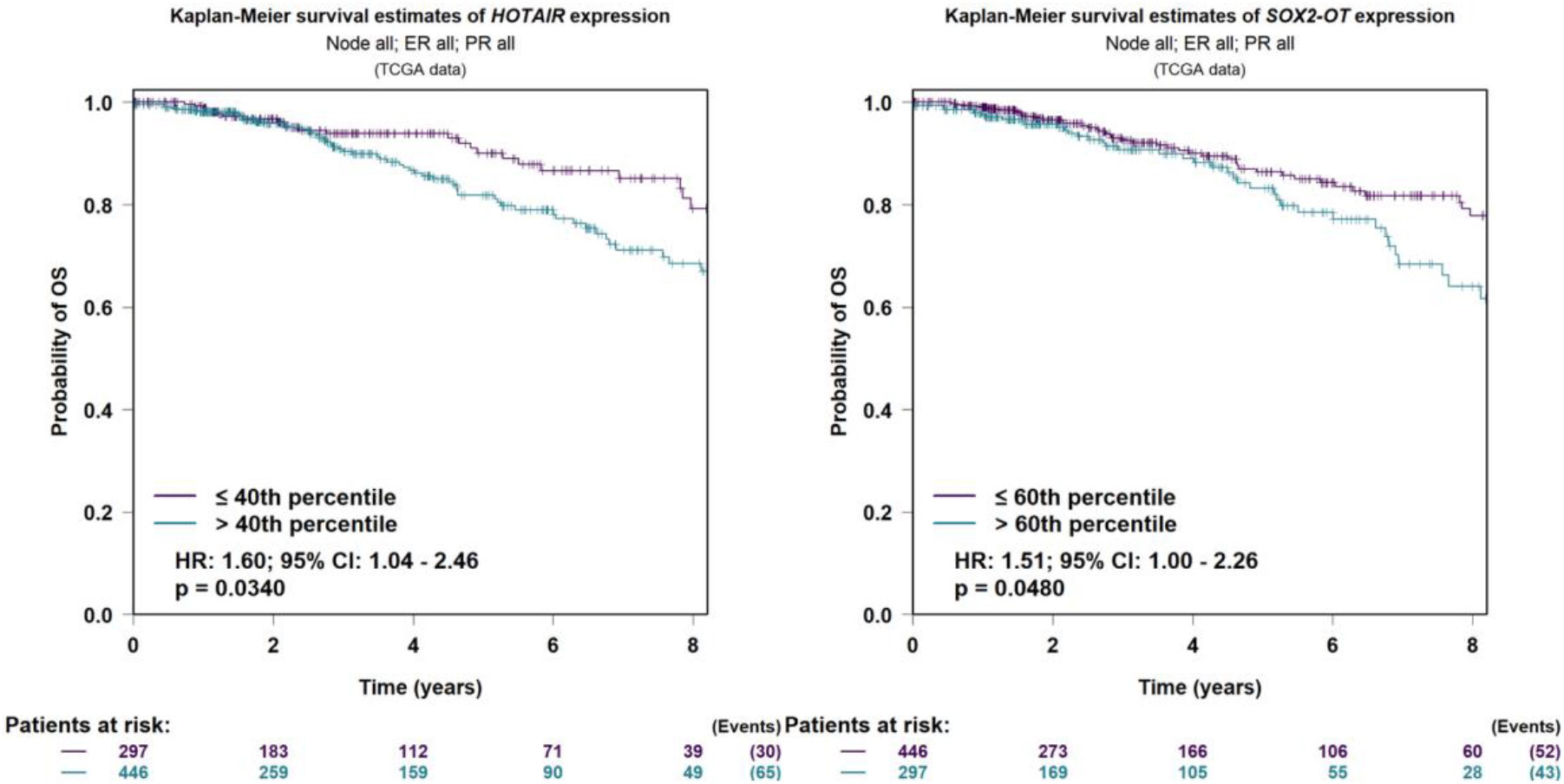
Survival analysis for the top predicted lncRNAs in breast cancer

## Discussion

The increasing body of evidence underscores the importance of developing a robust computational approach for elucidating potential associations between lncRNAs and disease, thereby facilitating an in-depth exploration of complex diseases at the molecular level. This work holds significant implications for advancing the diagnosis, treatment, prognosis, and prevention of various diseases, contributing valuable insights to the field of health science. Existing models for predicting lncRNA-disease associations often overlook the rich information embedded within nodes in the heterogeneous graph. Meta-paths, allowing the definition of diverse relationship paths between nodes of different types, offer a flexible means to capture the intricate structure and semantic information within heterogeneous graphs, while mitigating computational intricacies. Nevertheless, one problem with current meta-path-based lncRNA-disease association prediction models is that they frequently rely on pre-defined meta-paths based on substantial biological knowledge. To address the limitations associated with manually chosen meta-paths, we proposed the AMPGLDA model, capable of adaptively generating multiple meta-paths. Our evaluation demonstrated the superior performance of AMPGLDA compared to other LDA prediction models, as evidenced by higher AUC and AUPR values in 5-fold cross-validation. Additionally, the model’s robustness and generalization ability are confirmed across three distinct datasets. Finally, our investigation into three cancers substantiates the AMPGLDA model’s exceptional capability to uncover lncRNAs associated with underlying diseases.

The superior performance of AMPGLDA can be attributed to several factors. Firstly, we employed a devise range of biological data to construct a heterogeneous graph, resulting in a piece of comprehensive information. The heterogeneous graphs fused experimental support for LDAs and MDAs, functional similarity of lncRNA, semantic similarity of diseases, and interaction between lncRNA and miRNAs. Secondly, AMPGLDA mitigates the computational complexity of the model by adaptively extracting effective meta-paths, thereby flexibly capturing complex structures and semantic information within heterogeneous graphs and extracting potential feature representations. This differs from previous meta-path methods that rely on predefined biological knowledge. This approach avoids the redundancy and noise arising from arbitrary meta-path selection and addresses the issue of poor prediction performance associated with such choices. Thirdly, the inclusion of MLP enhances the model’s predictive capability. MLP effectively learns features from lncRNA-disease pairs, thereby improving the model’s fault tolerance and enhancing the efficiency and speed of prediction results.

Furthermore, our model comes with certain limitations:

1. There are a large number of lncRNA and diseases, but the associations verified by experiments are very small, which leads to the imbalance between the number of positive and negative samples in the process of model training. At present, the approach of randomly selecting negative samples poses a potential impediment to the efficacy of model training. The reason is that negative samples may potentially include positive samples that lack experimental verification but have an actual association, rendering them not truly negative samples. Hence, it is imperative to devise a more rational strategy for negative sample selection in future research.
2. Moreover, the model exhibited bias in calculating similarity, specifically in the computation of lncRNA functional similarity using known lncRNA-disease associations. Consequently, in subsequent research, we aim to explore more effective methods for calculating similarity to enhance the model’s performance.

## Conclusion

It is common knowledge that meta-path is crucial in capturing various semantic relationships between network components. The challenge of manually selecting the appropriate meta-path from a complex heterogeneous network has been effectively addressed by AMPGLDA through adaptive meta-path selection and integration with PCA for lncRNA-disease association prediction. Experimental results have indicated that the AUC and AUPR of AMPGLDA exceed other advanced LDA prediction methods. This breakthrough provides a fresh perspective on discovering disease-associated lncRNA. The confirmation of AMPGLDA’s predictive capability in diseases like breast, lung, and colorectal cancer through case studies further attests to its effectiveness. Furthermore, AMPGLDA holds the potential to be applied in diverse link prediction endeavors, encompassing miRNA-disease association prediction, circRNA-disease association forecasting, lncRNA-miRNA interactions, and more.

## Materials and methods

In this study, we propose a novel lncRNA-disease prediction method called AMPGLDA, as shown in **Figure 3**. Firstly, we introduced miRNA nodes, and leveraging biological information, constructed a lncRNA-disease-miRNA heterogeneous graph, capturing the complex interconnections between these three types of nodes. Global similarity feature representations of nodes were extracted using PCA to integrate information from different nodes and intricate edge junctions. Subsequently, the meta-path was learned adaptively from the heterogeneous graph, covering various meta-path lengths between lncRNA and disease. The features obtained from these two approaches were fused to create the final node feature representation. Ultimately, the lncRNA-disease association was scored using the MLP classifier.

**Figure 3.**
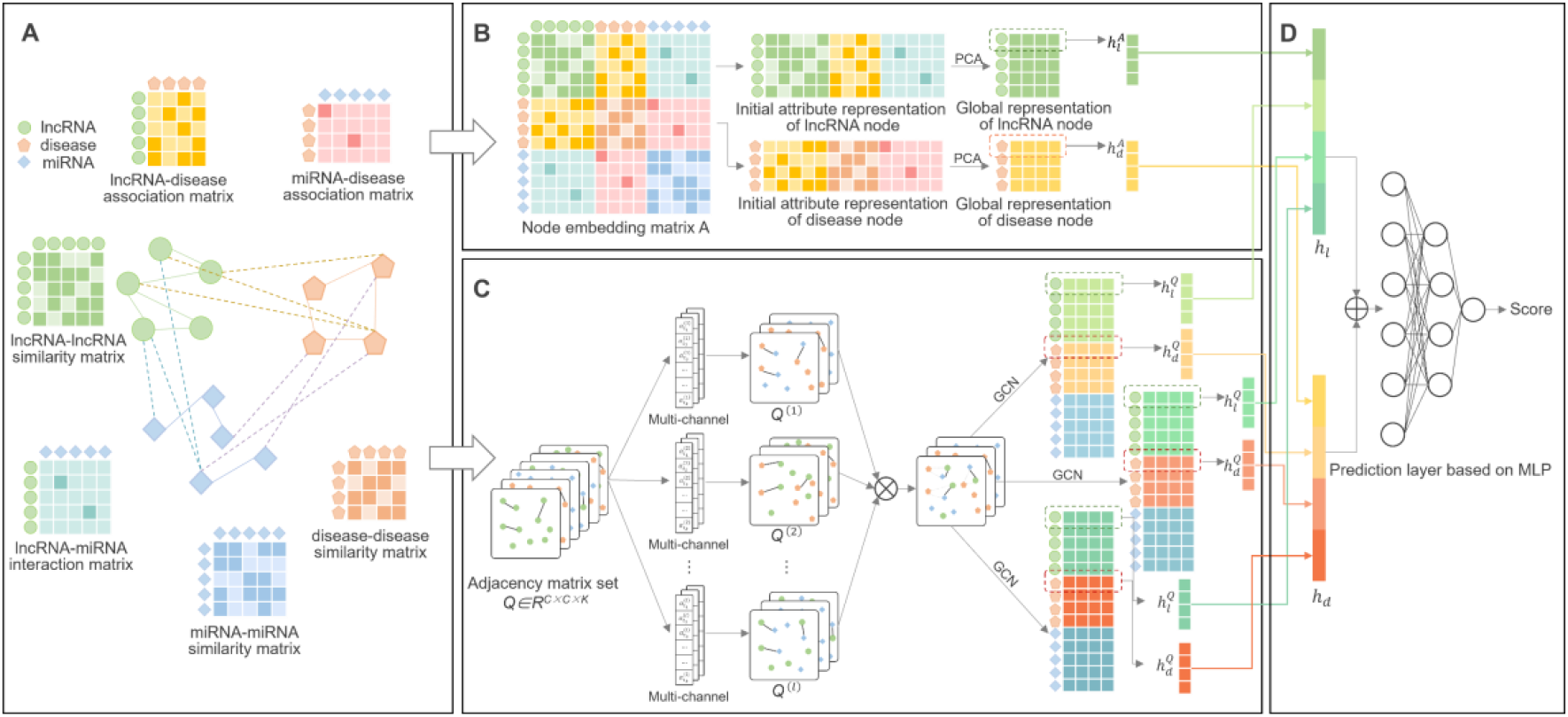
The flowchart of AMPGLDA. **A.** Construct heterogeneous network: construct a three-layer heterogeneous network using lncRNA, miRNA, and disease data. **B**. Node similarity features: the adjacency matrix A was obtained based on heterogeneous networks, and the similarity between lncRNA and disease was obtained by PCA dimensionality reduction. **C**. Node semantic feature: construct adjacency matrix set Q based on a heterogeneous network, and get the graph representation of meta-path by the meta-path adaptive learning method we designed, the semantic features of lncRNA and disease are represented by GCN layers. **D**. Feature fusion and prediction layer: lncRNA and disease similarity and semantic feature representation were performed for feature fusion to get the final feature representation, and lncRNA association score with disease was obtained by input into MLP.

### Datasets

The dataset utilized in this study, comprising 240 lncRNAs, 412 diseases, and 495 miRNAs, was sourced from a prior study by Fu et al. [20]. More specifically, the data were extracted from the Lnc2Cancer database [43], LncRNADisease database [44], and GeneRIF database [45], providing 2697 pairs of lncRNA-disease associations. Additionally, HMDD [46] contributed 13562 pairs of miRNA-disease associations, and starBase v2.0 [47] contributed 1002 pairs of miRNA and lncRNA interactions.

### Similarity calculation

#### Disease semantic similarity

This paper employs the approach of Wang et al. [48] to calculate the semantic similarity among various disease terms. Each disease corresponds to a node within a Directed Acyclic Graph (DAG) structure. The hierarchical connections among these nodes are leveraged to determine the semantic similarity between any pair of diseases. The semantic value of a particular disease node (denoted as “*d*”) is computed using the following formula (Equation 6):

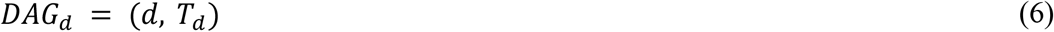

Where d denotes the disease node in the DAG, *T*_*d*_ denotes the union of edges connecting these nodes. The formula for determining the semantic contribution of each corresponding disease node is as follows:

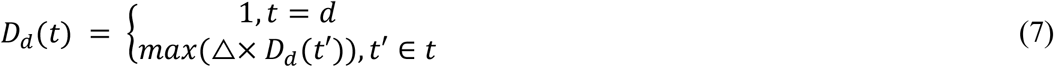

Here, Δ denotes the semantic contribution factor, which ranges from 0 to 1. In general, a value of 0.5 is more appropriate for Δ as it reflects the impact of the parent node on the child node within the DAG structure. The sum of semantic contribution values of the disease nodes associated with disease node “*d*” in the DAG structure is known as the semantic value (*DV*) of disease “*d*”. The following is the calculation formula:

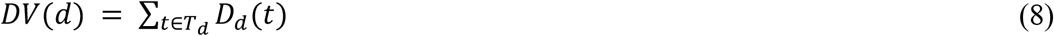

It is worth noting that the semantic similarity between two diseases, denoted as *d*_1_ and *d*_2_, exhibits a direct correlation with the quantity of shared nodes within the DAG structure. This similarity is calculated using the following formula:

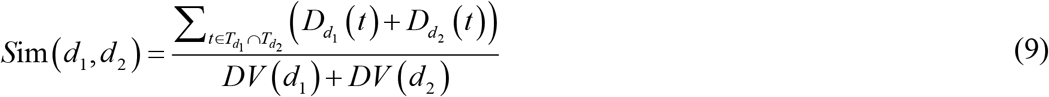

Where *t* denotes the common node of disease *d*_1_ and *d*_2_ in the DAG structure, the semantic values of disease *d*_1_ and *d*_2_ are represented by *DV*(*d*_1_) and *DV*(*d*_2_). 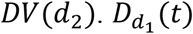 and 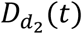 represent the semantic contributions of disease *t* to disease *d*_1_ and *d*_2_ , respectively. The semantic contributions of disease *t* to disease *d*_1_ and *d*_2_ are expressed as 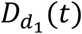 and 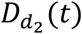, respectively.

#### lncRNA / miRNA Functional Similarity

Subsequently, we utilized the approach developed by Wang et al. [49] to establish the functional similarity between lncRNA, taking into account the previously calculated disease similarity and LDAs. For a given node of lncRNA / miRNA *r*_1_ and *r*_2_ , assuming that *n*_1_ diseases related to *r*_1_ and *n*_2_ diseases related to *r*_2_, denoted by *t*_1i_, (1≤i≤*n*_1_) and *t*_2j_, (1≤j≤*n*_2_), respectively, the functional similarity between *r*_1_ and *r*_2_ is :

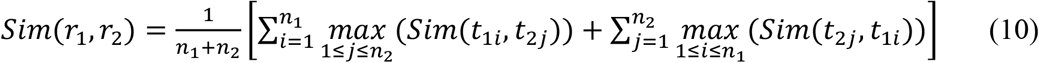

### Constructing a multi-view heterogeneous graph of lncRNA-disease-miRNA relationships

The input of our model is a multi-graph structure with multiple kinds of nodes and edges. In this structure, the association and similarity among nodes are regarded as the connection between nodes **(Figure 3A)**. Firstly, we establish a three-layer heterogeneous graph *G* = (*V, E*) using association and similarity information from two different perspectives. Three different types of nodes are included in the nodes set *V* = {*V*_*lnc*_⋃*V*_*dis*_⋃*V*_*mi*_}: lncRNA (*V*_*lnc*_), disease (*V*_*dis*_), and miRNA (*V*_*mi*_). The edge set *T*^*e*^ = {*A*_*l*−*d*_⋃*A*_*m*−*d*_⋃*A*_*l*−*m*_⋃*S*_*l*_⋃*S*_*m*_⋃*S*_*d*_} contains six lncRNA and miRNA-related networks: lncRNA-disease association set (*A*_*l*−*d*_ ∈ *R*^*l*∗*d*^), miRNA-disease association set ( *A*_*m*−*d*_ ∈ *R*^*m*∗*d*^ ), lncRNA-miRNA interaction set ( *A*_*l*−*m*_ ∈ *R*^*l*∗*m*^ ), lincRNA similarity (*S*_*l*_ ∈ *R*^*l*∗*l*^), miRNA similarity (*S*_*m*_ ∈ *R*^*m*∗*m*^), disease similarity (*S*_*d*_ ∈ *R*^*d*∗*d*^). We classified the lncRNA and miRNA nodes based on their associations with the disease node, setting the corresponding element in the adjacency matrix *A*_*ij*_ ∈ *A*_*l*−*d*_(*A*_*m*−*d*_ *or A*_*l*−*m*_) to 1 if they were associated (in the case of lncRNA, miRNA, and disease nodes) or interacted (for lncRNA-miRNA nodes). If there was no association or interaction, the element was set to 0.

### Global representation based on PCA

By combining the above information with the heterogeneous graph, we can obtain a complete weighted adjacency matrix by splicing:

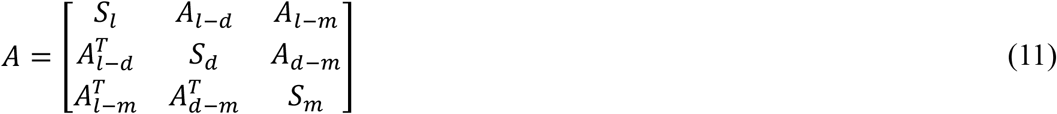

where 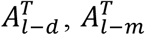, and 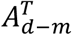 denote the transpose of *A*_*l*−*d*_ , *A*_*l*−*m*_,and *A*_*d*−*m*_. Therefore, using the acquired weighted adjacency matrix A, we construct global feature representations of nodes **(Figure 3B)**. The global representation of lncRNA can be thought of as the first l rows corresponding to lncRNA, and the global representation of disease nodes as the rows corresponding to disease. The low-dimensional embedding of lncRNA and disease was obtained by PCA.

### Semantic representation based on Meta-path generation

#### Meta-path generation adaptively

Previous studies [30] required manual pre-definition of meta-paths and combined deep learning to predict lncRNA-disease associations. This approach heavily relied on biological expertise, which severely limited the model’s scalability and flexibility and even impacted the model’s performance. To automatically discover more effective meta-paths, our method designs an adaptive meta-path learning module based on a graph transformer network [50]. This module automatically discovers, in heterogeneous graphs, meta-paths of any length between lncRNA and disease. These meta-paths are networks of paths joined by multiple-hop connections and heterogeneous edges **(Figure 3C)**.

Firstly, a set of adjacency matrix *Q* ∈ *R*^*C*×*C*×*K*^ is constructed according to the nodes and edge types in the three-layer heterogeneous graph, where *K* = |*T*^*e*^| and each adjacency matrix *Q*_*t*_ ∈ *R*^*C*×*C*^ . Here, C represents the number of nodes in the heterogeneous graph. For example, the adjacency matrix 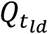 with the edge type of lncRNA-disease is expressed as 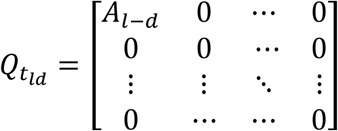, Other similar.

Then, the graph transfer layer softly selects two adjacency matrices from the adjacency matrix *Q* and learns a new graph structure by matrix multiplication of the two adjacency matrices. In order to realize the automatic generation of meta-paths, the graph transfer layer softly selects two adjacency matrices 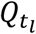 from *Q* through the convex combination of adjacency matrices (Formula 7).

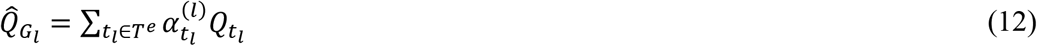

Where *T*^*e*^ is the set of edge types, 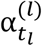 is the weight of the edge type *t*_l_ in the *l*th soft selection, and 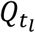 is the adjacency matrix of the *l*th layer.

The path graph is obtained by matrix multiplication of two adjacent matrices, and normalized to ensure numerical stability (Formula 8). For example, a learned meta-path 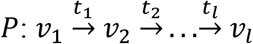,where *v*_*i*_ ∈ *V*,*t*_*i*_ ∈ *T*^*e*^, then the adjacency matrix *A*_*P*_ of the meta-path graph *G*_*p*_ automatically learned by *P* can be calculated by Formula 9.

To calculate the meta-path graph structure, given two soft selection adjacency matrices *Q*_1_ and *Q*_2_, we multiply them (*Q*_1_*Q*_2_) to obtain the meta-path graph structure, which is then normalized to ensure the numerical stability, that is,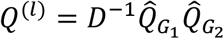. To learn multiple meta-paths, we use *l* channels to obtain multiple meta-path graphs. The calculation formula of the final adjacency matrix *A*_*P*_ of any length meta-path is as follows:

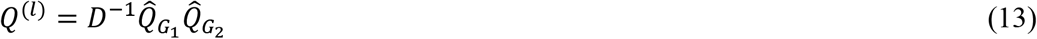

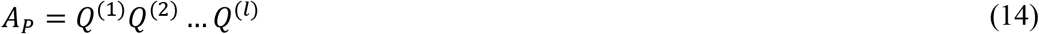

It is also worth mentioning that to learn the meta-path of any length containing the original edge, we add the identity matrix *A*_0_ = *I* to *Q*.

#### Learning node representations from multiple meta-path graphs

At present, GCN havs demonstrated efficacy in both theoretical understanding and practical applications for extracting graph structure features. GCN excels in integrating edge-related information of nodes, generating low-dimensional feature representations. The multiple meta-path graphs constructed earlier serve as inputs for GCN to aggregate the neighbor node information at each layer, along with semantic information from the edges of the meta-path. Consequently, this process yields enhanced representations for lncRNA and disease nodes. These refined representations are then fused with the global similarity features of the nodes, culminating in the acquisition of the final lncRNA and miRNA node embedding. The feature representation of the lth layer node in GCNs is defined as *H*^(*l*)^ , and the forward propagation relationship between layers is:

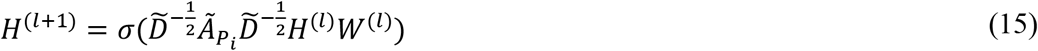

Where *σ* represents the nonlinear activation function, and *W*^(*l*)^ is a learnable weight matrix that graphs feature to latent space. 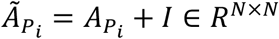 is the adjacency matrix of the graph, and 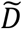 is the diagonal node degree matrix of 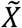.

### Final integration and prediction layer

Using a multi-layer convolution layer, we obtain feature representations of lncRNA and disease from the diverse meta-path graphs, denoted as 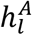 and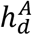, respectively. Next, to contain additional potential details in the feature representation of lncRNA and disease, we fuse 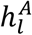 and 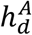 with the initial global features 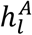 and 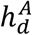 of lncRNA and disease, respectively. The most concise feature aggregation method usually uses the features of the two pathways in series to form a long vector, so the resulting lncRNA and disease features are represented as Formula 11 and Formula 12, respectively **(Figure 3D)**.

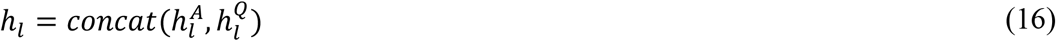

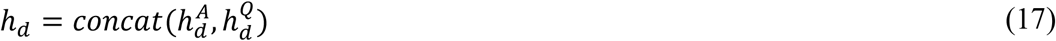

Among them, *h*_*l*_ represents the ultimate embedding for lncRNA, and the ultimate embedding of disease represents *h*_*d*_. In this representation, the embedding of each lncRNA can be represented by one row, and a similar representation is applied for diseases.

The embedding *h*_*l*_ of each lncRNA and the embedding *h*_*d*_ of each disease are spliced after obtaining the final embedding of lncRNA and disease to acquire our complete lncRNA-disease pairs embedding H, as shown below:

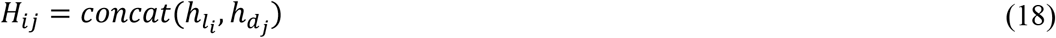

Where *H*_*ij*_ represents the characteristics of *l*_*i*_ and *d*_*j*_ in lncRNA-disease pairs. However, the aforementioned simplistic approach of feature fusion fails to capture the complex interplay between lncRNA and disease. To address this limitation, we propose combining the network embeddings of lncRNA and disease and leveraging MLP to make high-quality predictions for lncRNA-disease associations. Specifically, we employ the following equation to achieve our goal:

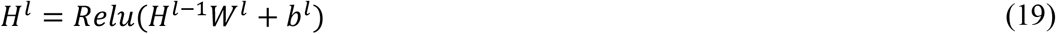

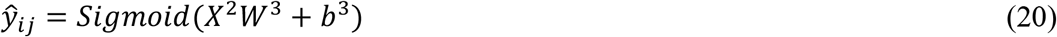

Here, *H*^*l*^ denotes the output of the first hidden layer, and *W*^*l*^ and *b*^*l*^ refer to the parameters and bias of that layer, respectively. We optimize our model by minimizing the error of the binary cross entropy loss function, as displayed in follows:

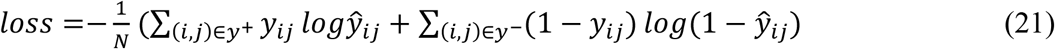

Where (*i, j*) represents the lncRNA-disease pair (*l*_*i*_, *d*_*j*_) , *y*^+^ and *y*^−^ represent positive and negative sample sets, and *N* represents the total number of lncRNA-disease pairs in such sets.

## CRediT author statement

**Xuehui Zhang:** Conceptualization, Investigation, Methodology, Writing – original draft, Validation. **Dengju Yao:** Supervision, Formal analysis, Writing – review & editing. **Xiaojuan Zhan:** Data curation, Writing – review & editing.

## Acknowledgments

This work is supported by the National Natural Science Foundation of China (Grant No. 62172128). The funding body did not play any roles in the design of the study and collection, analysis, and interpretation of data and in writing the manuscript.

## Declaration of AI and AI-assisted technologies in the writing process

In this paper, there is no reference to AI auxiliary technology in the writing process.

